# Computational Discovery of Core Protein–Targeted Therapeutics Against Ginger Wilt Pathogen *Ralstonia solanacearum*

**DOI:** 10.1101/2025.10.30.684991

**Authors:** Tianyue Zhang, Xiangyu Shen, Jiankang Tan, Jun Wang, Yiling Qin, Huansheng Cao, Nengfei Wang

## Abstract

Bacterial wilt caused by *Ralstonia solanacearum* poses a severe threat to ginger production worldwide, as no consistently effective control measures exist. Here, we present a comprehensive strategy integrating pangenome analysis, functional enrichment, molecular docking, and state-of-the-art protein structure prediction to identify antibacterial agents targeting this phytopathogen. By prioritizing core, evolutionarily conserved proteins in *R. solanacearum*, we utilized AutoDock Vina and the recently developed AlphaFold3 to assess the binding affinity and validate protein-ligand interactions of key candidates from commercially available antibacterial pesticides. In vitro antibacterial assays confirmed strong inhibitory effects for several compounds, notably streptomycin sulfate, phellodendrine chloride, chlorogenic acid, and allicin. Our results highlight the power of combining large-scale computational screening with experimental validation for accelerating the discovery of novel antibacterial agents against highly divergent, recalcitrant plant pathogens.

**Importance:** The indiscriminate use of broad-spectrum agrochemicals causes pollution and antimicrobial resistance. Our study proposes a targeted strategy against *Ralstonia solanacearum* by focusing on essential core proteins. This approach aims to develop specific agents that minimize harm to soil microbiota and reduce resistance selection. Our in silico framework aligns with green chemistry, offering a sustainable path for crop protection with a lower environmental footprint.

## Introduction

Ginger (Zingiber officinale) is a globally significant crop valued for its culinary and medicinal properties, but production has been severely threatened by bacterial wilt caused by *Ralstonia solanacearum*, especially at the critical stem–leaf junction. This soil-borne pathogen is notorious for causing rapid, destructive wilting, stunting, and rhizome rot, with some outbreaks resulting in total crop loss and catastrophic economic impacts for small-scale farmers and commercial producers alike. The bacterium infects its host through wounds and natural openings, colonizes the xylem, and forms resilient biofilms that block water transport, causing whole plants to wilt and die. Its virulence depends on cell wall-degrading enzymes and an arsenal of Type III secreted effectors that suppress host immunity, leading to widespread infection, compromised plant health, and reduced productivity. [1, 2]

Genomic studies reveal that *Ralstonia solanacearum* is highly adaptable, displaying extensive genetic diversity and possessing a complex genome with more than 5,000 coding sequences. Comparative and pangenomic analyses have identified hundreds of core and accessory genes, including over 70 Type III effectors, that account for its exceptional evolutionary potential and broad host range. Despite research into integrated and biocontrol management approaches, there are no consistently effective treatments against this pathogen; resistance breeding in ginger is limited by a lack of natural resistance sources, and chemical controls have proven inadequate due to the pathogen’s persistence and biofilm-mediated resilience. Persistent survival in soil and plant debris, ease of transmission, and genetic plasticity make *R. solanacearum* extremely hard to eliminate. [2, 3]

Recent advances in computational biology, particularly protein drug target identification using AutoDock and AlphaFold, are pushing the boundaries of rational drug development for recalcitrant phytopathogens. Molecular docking and machine learning-based protein structure prediction enable high-throughput virtual screening and precise mapping of protein-ligand interactions, as demonstrated in “one target, many compounds” approaches and benchmarking studies. These tools have shown predictive power for certain essential bacterial proteins and are being combined with structure-based drug discovery to identify candidate molecules that can block pathogen growth or virulence factors. In *Ralstonia solanacearum*, subtractive genome analysis and in silico target discovery have successfully pinpointed unique metabolic pathways and essential non-homologous proteins as druggable targets; docking simulations and virtual screening are now used to prioritize molecules for further validation. [4-6]

The aim of this study is to provide a new paradigm for identifying antibacterial agents against *Ralstonia solanacearum* by leveraging comprehensive pangenome analysis to systematically uncover core, accessory, and strain-specific drug targets, followed by molecular docking to screen and validate candidate compounds against these targets. This integrative approach, combining comparative genomics, protein structure prediction, and in silico ligand screening, offers a scalable framework for precision drug design that could overcome the limitations of conventional control methods and lay the foundation for sustainable disease management in ginger. Ultimately, molecular docking and target validation will allow not just identification, but also experimental confirmation of new drugs against validated protein targets, paving the way for robust disease control strategies. [7]

## Methods

### Bacterial Strain and Antibacterial Experiment

The test strain, *Ralstonia solanacearum*, was isolated from ginger and maintained in our laboratory. The bacterium was cultured overnight in LB (Luria-Bertani) broth (10 g/L tryptone, 5 g/L yeast extract, 10 g/L NaCl) at 30 °C with shaking at 150 rpm. Cells were diluted with fresh LB medium and grown to the logarithmic phase, corresponding to an optical density at 600 nm (OD_600_) of about 0.5 and approximately 1×10^9^ CFU/mL, as calibrated by serial dilution plating.

Four commercial antibacterial formulations, specifically marketed for the control of bacterial wilt and other bacterial diseases, were tested: zhongshengmycin, QingkuliKe, agricultural streptomycin, and garlic oil. The major active ingredients and their concentrations in each formulation are provided in Table 1. Stock solutions were prepared in sterile LB broth at the recommended field concentration, according to manufacturer instructions. For formulations that were insoluble or turbid, suspensions were treated with ultrasonic waves (300 W, 15 min, intermittent mode: 10 s on/10 s off) to ensure sterilization. The sterility of the processed solutions was confirmed by plating on LB agar before use.

**Table 1.** Information of four commercial antibacterials.

| Commercial Name | Main Active Ingredient | Physical Form | Recommended Concentration |
| --- | --- | --- | --- |
| Zhongshengmycin | Zhongshengmycin | Powder | 20 mg/mL -25 mg/mL |
| QingkuliKe | Astragalus Polysacharin | Liquid | 5 μL/mL |
|  | Chlorogenic Acid |  |  |
|  | Phellodendrine |  |  |
|  | Berberine |  |  |
| Agricultural Streptomycin | Streptomycin | Powder | 12.5 mg/mL-33 mg/mL |
| Garlic oil | Allicin | Liquid | 1 μL /mL |

Working solutions were prepared by two-fold serial dilution, with the recommended concentration as the midpoint, and stepwise dilutions above and below. For each treatment, 100 µL of bacterial suspension (∼1×10^9^ CFU/mL) was added to 10 mL of antibacterial solution. Cultures were incubated at 30 °C under static conditions for 24 h. Controls included antibacterial solution without bacteria and bacterial suspension without antibacterial.

After incubation, 100 µL aliquots from each treatment were spread on TTC agar plates and incubated at 30 °C for 48 h. The minimum bactericidal concentration (MBC) was defined as the lowest antibacterial concentration that completely inhibited colony growth. Colonies of *Ralstonia solanacearum* on TTC agar were identified by the characteristic red coloration, which is a distinguishing feature of this bacterium. Each treatment included three technical replicates, and the entire experiment was independently repeated three times to ensure reproducibility.

### Selection of Docking Targets

To identify conserved receptor candidates, 208 genomes of *Ralstonia solanacearum* were retrieved from the NCBI RefSeq and GenBank databases. Only high-quality assemblies (scaffold, chromosome, or complete level) were included. Genome annotation was standardized, and pangenome construction was performed at a 95% BLASTp identity threshold. Genes were categorized into four groups: core, soft-core, shell, and cloud, with core genes defined as those present in ≥98% of strains.

To ensure functional relevance and structural reliability, genes annotated as “putative proteins,” “uncharacterized proteins,” “putative transport proteins,” “conserved putative proteins,” fragments or partial proteins, short peptides, mobile element proteins, and phage proteins were systematically excluded. The remaining core genes were mapped to the UniProt database, and only those with corresponding protein products available as crystal structures or high-confidence AlphaFold models (mean pLDDT ≥ 70) were retained for downstream analysis.

Functional prioritization of the filtered core genes was conducted using Gene Ontology (GO) enrichment analysis and KEGG pathway enrichment analysis. All filtered core genes were used as the background for GO enrichment, with significance assessed by Fisher’s exact test at *p* < 0.001. Enrichment consistency across multiple presence thresholds was used as an additional robustness check. Redundant GO terms were simplified using semantic similarity measures, and functional networks were visualized. KEGG pathways with adjusted *p*-values below 0.05 were considered statistically significant, and the top 20 pathways were selected based on P-value ranking.

Genes associated with significantly enriched GO terms or the top 20 KEGG pathways were identified, and their corresponding protein products were retained as core proteins for downstream docking analysis. Duplicates appearing in both GO and KEGG sets were included only once. This approach provided a curated set of structurally characterized, evolutionarily conserved, and functionally relevant proteins suitable for ligand-receptor predictions.

### Molecular Docking

Three-dimensional receptor structures were preprocessed using MGLTools. Water molecules were removed, polar hydrogens added, and Gasteiger charges assigned before conversion into PDBQT format.

Ligands, including astragalus polysaccharide, berberine, chlorogenic acid, phellodendrine, streptomycin and zhongshengmycin, were retrieved from PubChem in SDF format. Structures were energy-minimized using the MMFF94 force field and converted to PDBQT with Galaxy tools.

Docking simulations were performed with AutoDock Vina. The exhaustiveness parameter was set to 8, and eight binding modes were generated for each receptor–ligand pair. Each docking experiment was conducted in triplicate to minimize stochastic variability. Conformations with binding affinities ≤ –7.0 kcal/mol, a widely adopted threshold for significant ligand–protein interactions and clustering RMSD ≤ 2.0 Å were considered reliable binding poses. [8] Protein– ligand interactions were visualized with PyMOL.

### Assessment of Docking Predictions Using AlphaFold3

To provide a complementary evaluation of the docking results, AlphaFold3 was employed for structural modeling of selected protein–ligand complexes. Specifically, the three protein–ligand pairs with the highest predicted binding affinities from the AutoDock Vina calculations were chosen for further analysis. For each target, the corresponding protein amino acid sequence and ligand SMILES string were prepared and submitted to AlphaFold3.

The AlphaFold3-predicted complexes were compared with the docking poses obtained from AutoDock Vina, focusing on the consistency of binding site location, binding mode, and key interacting residues. The quality of the predicted complexes was evaluated using the predicted TM-score (ptm) and the interface predicted TM-score (iptm) provided by AlphaFold3. Following recommended guidelines, predictions with ptm > 0.5 and iptm > 0.8 were considered to be of very high confidence and were used in the comparative analysis. [9]

#### *In vitro* Validation

To experimentally assess the reliability of the molecular docking and AlphaFold3 predictions, we performed antibacterial assays using high-purity (≥98%) standards of the ligands identified as high-confidence binders. The selected compounds included astragalus polysaccharide, berberine hydrochloride, chlorogenic acid, phellodendrine hydrochloride, streptomycin sulfate, and allicin.

Zhongshengmycin was not included in the validation because a high-purity compound was not available. Assays were conducted in minimal M9 medium (composition detailed in Supplementary Table S1), chosen to mimic the nutrient-limited in planta environment.

Bacterial cultures at mid-exponential phase were inoculated into M9 medium supplemented with serial dilutions of each compound. For all compounds except allicin, growth was monitored by OD_600_ measurements during the first 6 h, capturing early growth dynamics before compound-induced sedimentation interfered with turbidity-based readings. At 12 h, colony-forming units (CFUs) were quantified by plating serial dilutions to provide an endpoint viability measure.

All solutions (except allicin) were sterilized by filtration through 0.22 μm membranes. Allicin was prepared as a stock solution in anhydrous ethanol; due to its oily nature, allicin stock formed emulsions upon dilution, precluding reliable OD-based growth monitoring. Therefore, the antibacterial activity of allicin was assessed exclusively by CFU enumeration. Solvent controls containing 5% ethanol were included in parallel to account for ethanol-derived effects.

## Results

### Preliminary Antibacterials Screening Results

After extensive search in literature and antibacterial pesticides against plant pathogens, four commercial antibacterial pesticides on the market were obtained. Initial screening of these four antibacterials for controlling bacterial wilt showed that Zhongshengmycin, QingkuliKe, and Garlic Oil exhibited clear Minimum Bactericidal Concentrations (MBCs) of 80 mg/mL, 500 μL/mL, and 125 μL/mL in treating *R. solanacearum*, respectively. In contrast, Agricultural Streptomycin did not yield a measurable MBC under the conditions tested. This lack of detection can be attributed to its relatively low effective concentration and the limited solubility of the formulation.

### Selection of Docking Targets

Next, we set out to identify the targets of these antibacterial pesticides in the pathogen *R. solanacearum*. To narrow down the possible targets of these antibacterial pesticides, which should be present in all or most *R. solanacearum* strains. Following this line of inference, we first identified the core proteins shared by *R. solanacearum* strains through pangenomic analysis using Roary. Results showed that 786 genes were present in at least 205 of the 208 analyzed genomes. The gene frequency distribution exhibited steep peaks on the left, confirming the observation that only small proportion of genes shared by all strains; and the majority of genes are present in only a small number of strains, consistent with the prevalence of strain-specific (cloud) genes. So now we had 786 genes as potential targets.

### Protein-compound pairs revealed by Molecular Docking

We further narrowed down the genes by performing functional enrichment of the gene products through GO enrichment analysis (Supplementary Table S2) and KEGG pathway analysis (Fig. 1C). The gene products must be associated with consistently enriched GO terms and KEGG pathways. A total of 1,562 proteins encoded by 786 core genes (Supplementary Table S3) were selected as targets. We also identified the effective components in antibacterial pesticides against the pathogen as ligands for molecular docking (Table 2). As shown in Fig. 2A, AutoDock results indicated that, except for the ligand allicin, all other ligands exhibited substantial predicted binding potential with the core proteins. Among these, berberine and chlorogenic acid displayed the highest median binding affinities (≤ –8 kcal/mol), whereas astragalus polysaccharide showed the lowest median binding affinity (slightly below –7 kcal/mol).

**Table 2.** Top molecular docking results from AutoDock and AlphaFold3.

| ligand | Receptor | affinity_kcal/mol | rmsd_lb | rmsd_ub | ptm | iptm |
| --- | --- | --- | --- | --- | --- | --- |
| Astragalus polysacharin | A0A0S4TUD0 | -8 | 0 | 0 | 0.75 | 0.77 |
|  | A0A0S4TZC3 | -7.9 | 0 | 0 | 0.94 | 0.82 |
|  | A0A0S4UZQ2 | -7.8 | 0 | 0 | 0.89 | 0.74 |
| Berberine | A0A0S4X5D9 | -10.3 | 0 | 0 | 0.87 | 0.75 |
|  | A0A0S4TW40 | -10.2 | 0 | 0 | 0.93 | 0.87 |
|  | A0A0S4WS66 | -10 | 0 | 0 | 0.82 | 0.87 |
| Chlorogenic acid | A0A0S4UBE2 | -10.5 | 0 | 0 | 0.9 | 0.75 |
|  | A0A0S4TZR9 | -10.3 | 0 | 0 | 0.92 | 0.65 |
|  | A0A0S4TZC3 | -10.1 | 0 | 0 | 0.94 | 0.8 |
| Phellodendrine | A0A0S4VXE3 | -9.3 | 0 | 0 | 0.63 | 0.58 |
|  | A0A0S4X6Q4 | -9.1 | 0 | 0 | 0.78 | 0.76 |
|  | A0A0S4U1N7 | -9 | 0 | 0 | 0.87 | 0.72 |
| Streptomycin | A0A0S4TW40 | -10.1 | 0 | 0 | 0.9 | 0.76 |
|  | A0A0S4U629 | -9.9 | 0 | 0 | 0.8 | 0.71 |
|  | A0A0S4TSY0 | -9.8 | 0 | 0 | 0.91 | 0.81 |
| Zhongshengmycin | A0A0S4TW40 | -9.7 | 0 | 0 | 0.91 | 0.78 |
|  | A0A0S4WPJ7 | -9.6 | 0 | 0 | 0.9 | 0.74 |
|  | G8GT43 | -9.3 | 0 | 0 | 0.76 | 0.54 |

**Figure 1.**
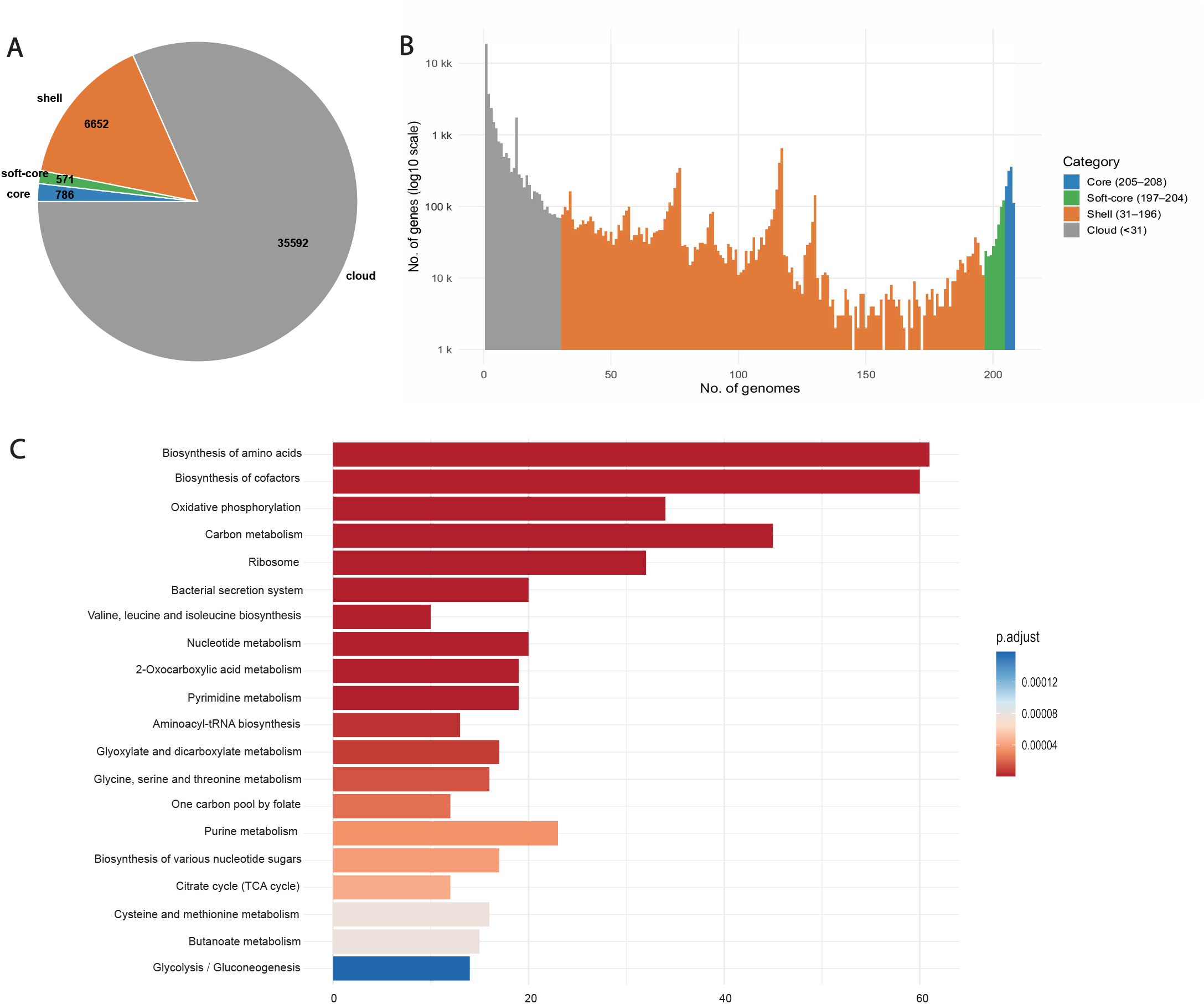
Pan-genome analysis of 208 *Ralstonia solanacearum* genomes and KEGG pathway enrichment of core genes. (A) Distribution of gene categories across the pan-genome. Core genes are present in 205–208 strains, soft-core genes in 197–204 strains, shell genes in 31–196 strains, and cloud genes in fewer than 31 strains. (B) Gene frequency distribution across genomes. Steep peaks on the left indicate that most genes are present in a minimal number of strains, highlighting the abundance of strain-specific (cloud) genes, whereas peaks on the right represent highly conserved core genes. (C) KEGG pathway enrichment analysis of core genes.

**Figure 2.**
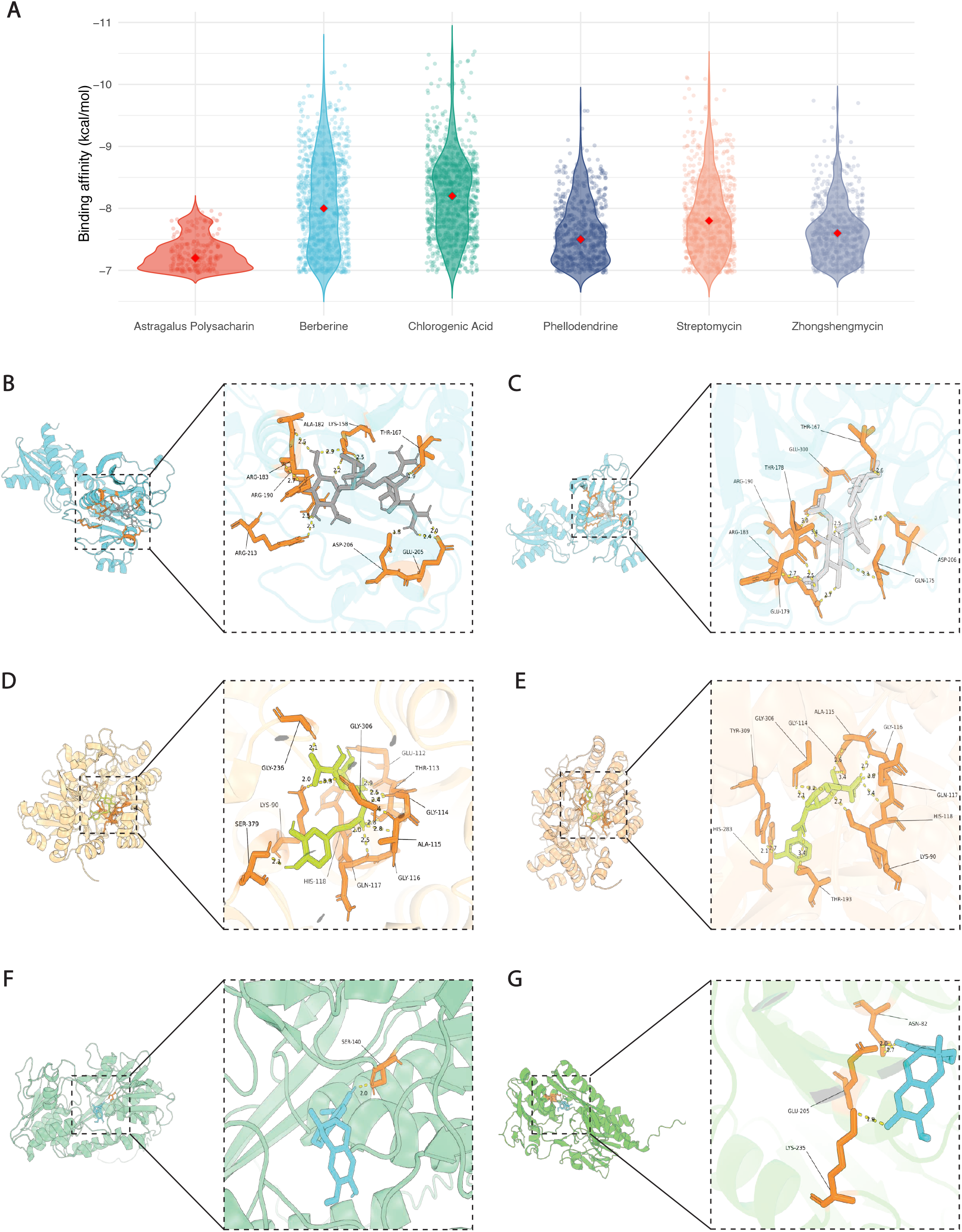
Docking and validation of ligand–core protein interactions. (A) Violin plots showing the distribution of binding affinities (≤ –7 kcal/mol) between six candidate ligands and core proteins, illustrating the overall variation in docking scores. (B, C) Predicted binding of streptomycin with protein A0A0S4TSY0 obtained by AutoDock (B) and by AlphaFold3 (C). (D, E) Predicted binding of chlorogenic acid with protein A0A0S4TZC3 obtained by AutoDock (D) and by AlphaFold3 (E). (F, G) Predicted binding of phellodendrine with protein A0A0S4U1N7 obtained by AutoDock (F) and by AlphaFold3 (G).

We further validated these binding affinities in silico using AlphaFold3. For each group, the top three ligand–protein pairs with the best predicted binding affinities were selected for validation using AlphaFold3, and the results are summarized in Table 2. Overall, the binding predictions from AutoDock and AlphaFold3 were highly correlated. Notably, in the phellodendrine and zhongshengmycin groups, one ligand–protein pair in each group showed a discrepancy, in which case AutoDock predicted strong binding while AF3 indicated lower binding potential.

We examined the specific bond types among the top protein-antibacterial pairs. First, consistent bond patterns were observed between AutoDock and AlphaFold3 predictions (Fig. 2B–G), with additional visualized docking results provided in the Supplementary Figure S1. For the interaction between streptomycin and protein A0A0S4TSY0, both approaches identified hydrogen-bond contacts with eight amino acid residues, and four of these residues (Thr167, Arg183, Arg190, and Asp206) were consistently predicted by both methods. For chlorogenic acid binding to protein A0A0S4TZC3, AutoDock predicted hydrogen-bond interactions with 11 residues, while AlphaFold3 predicted 10, with seven residues in common, including Lys90, Gly114, Ala115, Gly116, Gln117, His118, and Gly306. For phellodendrine binding to protein A0A0S4U1N7, AutoDock predicted a hydrogen-bond interaction with Ser140, whereas AlphaFold3 predicted interactions with Asn82, Glu205, and Lys235.

### *In Vitro* Validation of Docking Results

The predicted binding between protein targets and the effective components of the antibacterial pesticides were experimentally validated with the pathogen *R. solanacearum*. Based on the solubility properties of the effective compounds and pilot studies, the test concentrations were set as follows: astragalus polysaccharide at 1 mg/mL, streptomycin sulfate at 20 mg/mL, phellodendrine chloride at 5 mg/mL, berberine hydrochloride at 10 mg/mL, chlorogenic acid at 10 mg/mL, and allicin at 20 mg/mL. Preliminary OD_600_ growth curves indicated that astragalus polysaccharide promoted the growth of *R. solanacearum*, with growth rates higher than those of the control, and was therefore excluded from subsequent antibacterial assays. According to the regression analysis between OD_600_ values and colony-forming units (CFU) (Fig. 3A), the bacterial suspension used for inoculation at the beginning of the assay corresponded to the mid-exponential phase, with OD_600_ values of approximately 0.5, equivalent to a cell density of ∼10^9^ CFU/mL.

**Figure 3.**
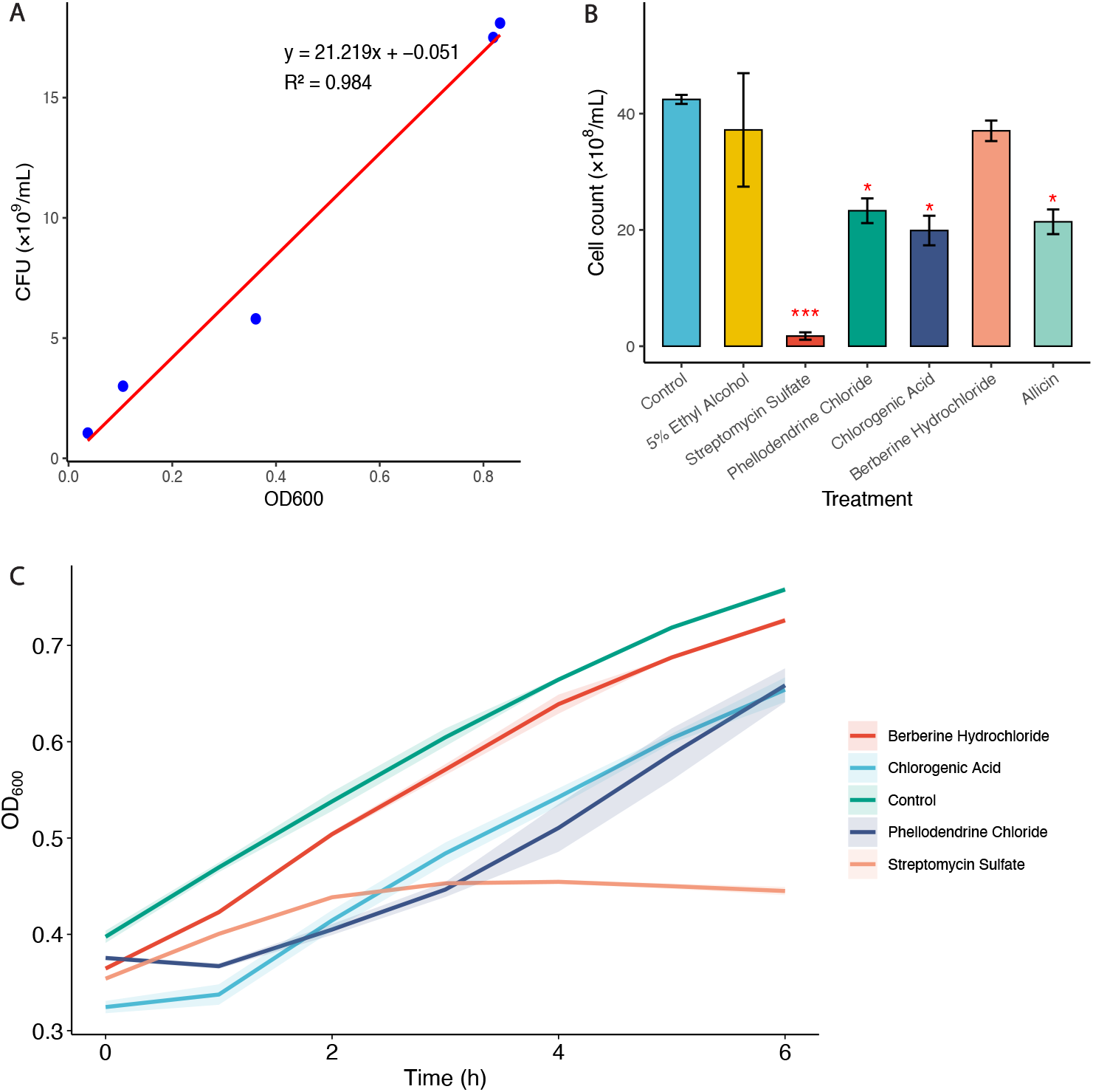
Growth dynamics and bactericidal effects of candidate compounds. (A) Regression curve between OD_600_ values and CFU counts in M9 medium. (B) OD_600_ growth curves of *Ralstonia solanacearum* over the first 6 h in the presence of different compounds, excluding Allicin, which was omitted because it is not water-soluble and would interfere with OD measurements. (C) Viable cell counts of each treatment group after 12 h of *in vitro* culture. Statistical significance was determined relative to the control group, with P < 0.05 (*), P < 0.01 (**), and P < 0.001 (***).

Figure 3.B shows the growth dynamics of *R. solanacearum* during the first six hours following the addition of effective antibacterial components. Compared with the control, the growth of the streptomycin sulfate treated group was markedly inhibited, with a noticeable in growth rate after 2 hours and a decline in cell density beginning at 4 hours. Phellodendrine chloride treatment resulted in an initial decrease in cell density, followed by gradual recovery, but the final OD values remained lower than those of the control. Chlorogenic acid treatment led to lower initial turbidity and slower early growth, which partially recovered after 1 hour, though overall growth remained below the control. Berberine hydrochloride-treated cells showed growth curves similar to the control.

To further assess antibacterial effects, viable cell counts were determined using the dilution plating method after 12 hours of culture, when *R. solanacearum* had reached the stationary phase (Fig. 3C). The 5% ethanol treatment, used as a solvent control for allicin, caused a slight reduction in colony numbers but showed no significant difference from the control. Quantitative analysis revealed that streptomycin sulfate reduced viable cell counts by 95.88% compared with the control (P = 0.041), representing the most pronounced effect among all treatments. Phellodendrine chloride, chlorogenic acid, and allicin treatments resulted in reductions of 45.11% (P = 0.015), 53.12% (P = 0.018), and 49.59% (P = 0.013), respectively. In contrast, berberine hydrochloride and 5% ethanol treatments showed only modest decreases of approximately 12% relative to the control, which were not statistically significant (P > 0.05).

## Discussion

This study demonstrates a new high-throughput approach for antibacterial discovery in plant pathogens by integrating molecular docking with AutoDock and the latest AlphaFold3 protein structure prediction platform. The combination enables the in silico screening of hundreds of core, functionally enriched protein targets, significantly accelerating the identification and prioritization of candidate small molecules with strong binding affinities. Notably, AlphaFold3’s diffusion-based architecture not only provides highly accurate protein structures but also enables detailed modeling of protein-ligand interactions, surpassing traditional approaches in prediction accuracy and speed, and offering an unprecedented opportunity for computationally driven drug discovery in phytopathology. [10-12]

Analysis of *R. solanacearum*’s pangenome revealed pronounced genetic divergence among strains, with a large shell and cloud genome and only a fraction of highly conserved core genes. These core targets, enriched in essential biological processes—including translation, energy metabolism, cell wall biosynthesis, and regulatory functions—represent the most promising nodes for chemical intervention that can impact pathogen fitness across diverse isolates. High divergence and rapid evolution among pathogenic lineages mean that broad-spectrum durability requires targeting such evolutionarily constrained proteins rather than accessory or strain-specific genes, which can rapidly be gained or lost. [13, 14] Functional characterization of the chosen targets reveals their critical roles in metabolic pathways, pathogenicity, and intrinsic resistance, underscoring their biological relevance. Our study demonstrates that virtual screening must be paired with robust experimental validation: most computational hits were supported by antibacterial assays, while false positives were efficiently excluded, highlighting the necessity of closed-loop, computational–experimental workflows. This integrative strategy is especially crucial for plant pathogens with high genomic plasticity and broad host range, such as *R. solanacearum*, because it enables iterative refinement and functional confirmation of predictions, ensuring that only biologically active compounds progress to field testing. [5, 12, 15, 16]

Overall, the application of advanced tools like AlphaFold3 and AutoDock, alongside rigorous in vitro validation, represents a new paradigm in rational antimicrobial discovery. This approach holds the potential to bridge the gap between computational predictions and practical disease management solutions, particularly in the face of emerging resistance and the dynamic evolution of plant pathogenic bacteria. [10, 11]

## Acknowledgements

This work was supported by National Natural Science Foundation of China (32171565), Duke Kunshan University Chancellor’s Fund, Kunshan Municipal Fund, and DKU-Duke Fund.

