## Supplementary Figure S1 for "Computational Discovery of Core Protein–Targeted Therapeutics Against Ginger Wilt Pathogen *Ralstonia solanacearum*"

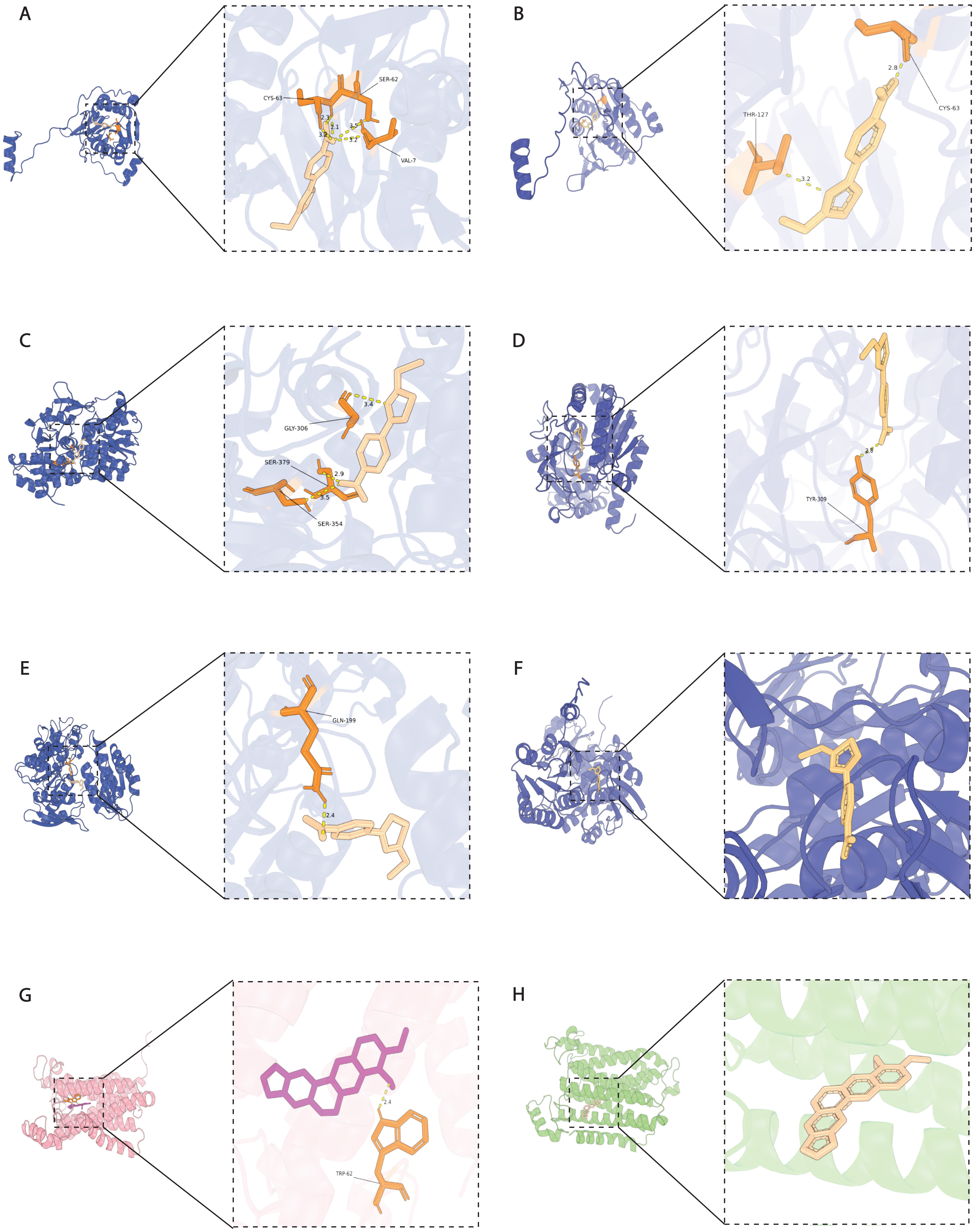
A
CYS-63
SER-62
VAL-7
2.3
2.1
3.5
3.2
B
2.8
CYS-63
THR-127
3.2
C
3.4
GLY-306
SER-379
2.9
3.5
SER-354
D
2.8
TYR-309
E
GLN-199
2.4
F
G
2.4
TRP-62
H

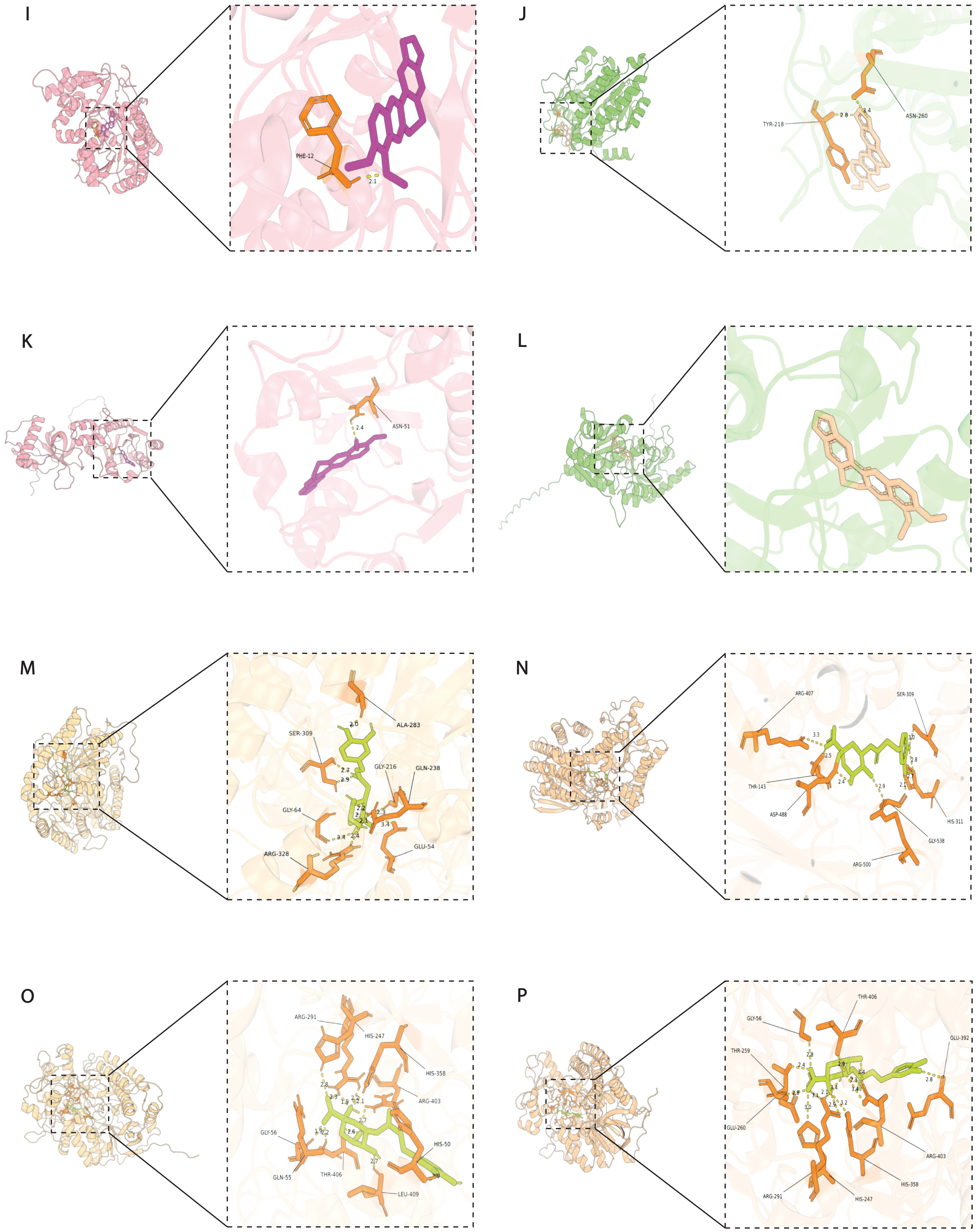
I
PHE-12
2.1
J
TYR-218
2.8
3.4
ASN-260
K
ASN-51
2.4
L
M
ALA-283
SER-309
GLY-216
GLN-238
GLY-64
ARG-328
GLU-54
N
ARG-407
SER-309
THR-143
ASP-488
HIS-311
GLY-538
ARG-500
O
ARG-291
HIS-247
HIS-358
ARG-403
GLY-56
HIS-50
GLN-55
THR-406
LEU-409
P
THR-406
GLY-56
GLU-392
THR-259
GLU-260
ARG-403
HIS-358
ARG-291
HIS-247

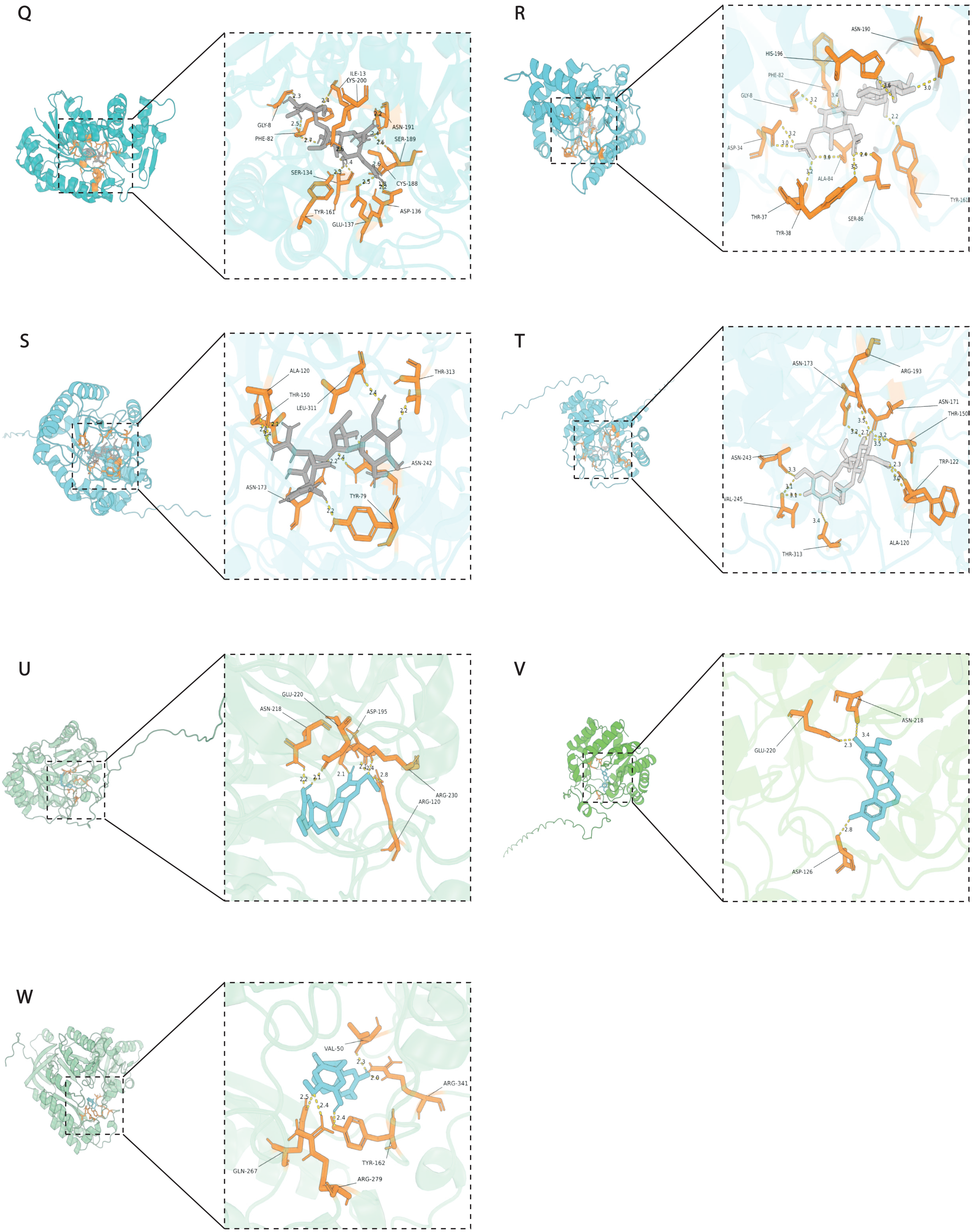
Q
ILE-13
LYS-200
GLY-8
PHE-82
ASN-191
SER-189
SER-134
CYS-188
TYR-161
ASP-136
GLU-137
R
ASN-190
HIS-196
PHE-82
GLY-8
ASP-34
ALA-84
TYR-161
THR-37
TYR-38
SER-86
S
ALA-120
THR-313
THR-150
LEU-311
ASN-242
ASN-173
TYR-79
T
ASN-173
ARG-193
ASN-171
THR-150
ASN-243
TRP-122
VAL-245
ALA-120
THR-313
U
GLU-220
ASN-218
ASP-195
ARG-230
ARG-120
V
ASN-218
GLU-220
ASP-126
W
VAL-50
ARG-341
TYR-162
GLN-267
ARG-279

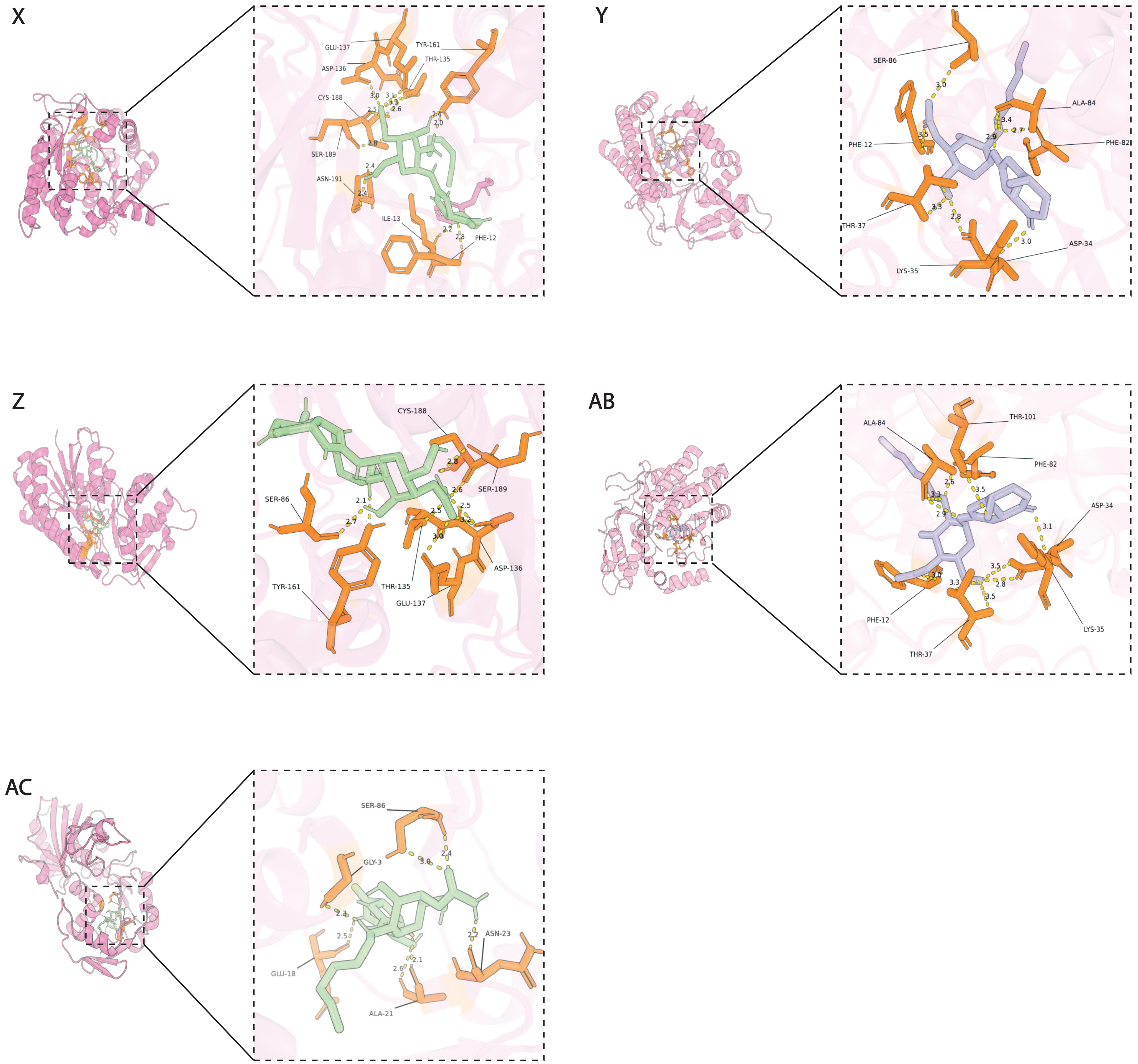


**Supplementary Figure 1**. Predicted binding of active compounds to target proteins. For each ligand–protein pair, left panels (A, C, E, ...) show Autodock predictions, and right panels (B, D, F, ...) show AlphaFold3 predictions. Alphafold3 predictions were unreliable for W and AC. Ligand–protein pairs: Astragalus Polysaccharin: A0A0S4TUD0 (A–B), A0A0S4TZC3 (C–D), A0A0S4UZQ2 (E–F); Berberine: A0A0S4X5D9 (G–H), A0A0S4TW40 (I–J), A0A0S4WS66 (K–L); Chlorogenic Acid: A0A0S4UBE2 (M–N), A0A0S4TZR9 (O–P); Streptomycin: A0A0S4TW40 (Q–R), A0A0S4U629 (S–T); Phellodendrine: A0A0S4X6Q4 (U–V), A0A0S4VXE3 (W); Zhongshengmycin: A0A0S4TW40 (X–Y), A0A0S4WPJ7 (Z–AB), G8GT43 (AC).
